# Staying in touch: how highly specialised moth pollinators track host plant phenology in unpredictable climates

**DOI:** 10.1101/2021.03.31.437762

**Authors:** Jonathan T. D. Finch, Sally A. Power, Justin A. Welbergen, James M. Cook

## Abstract

For pollinating insects that visit just a single flowering species, the co-occurrence of flowers and insects in time is likely to have critical implications for both plant and pollinator. Insects often utilise diapause to persist through periods in which resources are unavailable, timing their re-emergence by responding to the same environmental cues as their host plants. The obligate pollination mutualisms (OPMs) between *Epicephala* moths (Gracillariidae) and their leaf flower host plants are some of the most specialised interactions between plants and insects. However, to date there have been very few studies of *Epicephala* moth lifecycles and none of how they synchronise their activity with the flowering of their host plants. *Breynia oblongifolia* (Phyllanthaceae) is known to be exclusively pollinated by two highly specific species of *Epicephala* moth (Gracillariidae). We surveyed populations of both the host plant and it’s pollinators over multiple years to determine their annual phenology and then modelled the climatic factors that drive their activity. Using our newly gained knowledge of moth and host plant phenology, we then looked for evidence of diapause at both the egg and pre-pupal stages. Our phenology surveys showed that although female flowers were present throughout the entire year, the abundance of flowers and fruits was highly variable between sites and strongly associated with local rainfall and photoperiod. Fruit abundance, but not flower abundance, was a significant predictor of adult *Epicephala* activity, suggesting that eggs or early instar larvae diapause within dormant flowers and emerge as fruits mature. Searches of overwintering flowers confirmed this, with many containing evidence of pollen and diapausing pollinators. We also observed the behaviour of adult *Epicephala* prior to pupation and found that ~10% of the Autumn emerging *Epicephala* enter diapause, eclosing to adulthood after 38-56 weeks. The remaining 90% of autumn emerging adults pupate directly with no diapause, suggesting a bet hedging strategy for adult emergence. As such, *Epicephala* moths appear to utilise diapause at multiple stages in their lifecycle, and possibly bet hedging, in order to deal with variable flowering phenology and climatic unpredictability.

## Background

In ecosystems, resources are often ephemeral and unpredictable. How species track ephemeral resources, such as prey, fruits or flowers, is an important question in ecology (Armstrong *et al.* 2016; Deacy *et al.* 2016; Rivrud *et al.* 2018). Flowering can be influenced by variety of climatic factors including, temperature, rainfall and photoperiod (Davies 1976; Friedel *et al.* 1994; Jolly and Running 2004). The timing and intensity of some climatic factors, however, can be highly variable between years, making the distribution and occurrence of flowering resources unpredictable.

Pollinating insects that rely on a small number of flowering species, so-called specialists or oligotrophs, may be at greater risk of extinction due to a lack of available flowers (Encinas-Viso *et al.* 2012). Obligate pollination mutualisms (OPMs) are perhaps the most specialised interactions known to occur between plants and insect pollinators. In OPMs, insect pollinators generally transport pollen between the male and female flowers of a single host plant species. Along with pollen, female pollinators also deposit their eggs into female flowers. The ovules of the developing fruit then become the nursery and primary food source for the pollinator’s offspring. Many forms of OPM are currently known, the most widely studied being those occurring in figs (Cook and Rasplus 2003), *Yucca* (Pellmyr 2003), globeflowers (Thompson and Pellmyr 1992) and some members of the Phyllanthaceae family (Kawakita 2010). The OPMs occurring within the Phyllanthaceae or “leaf flowers” are the most recently discovered (~15ya) of the major OPM radiations (Kato *et al.* 2003), and it is now believed that up to 700 species of the genera *Breynia, Glochidion* and *Phyllanthus* are pollinated exclusively by *Epicephala* moths (Gracillariidae), also known as leaf flower moths (Kawakita and Kato 2009; Kawakita *et al.* 2019). As pollinators in OPMs are entirely reliant on the flowers of their host plant, the synchrony of plant and pollinator life history is critical to the persistence of pollinator populations and the stability of the mutualism (Bronstein *et al.* 1990).

Within OPMs there is a broad spectrum of flowering activity. OPMs in the tropics can flower continuously or near continuously (Bronstein *et al.* 1990; Kawakita and Kato 2004; Zhang *et al.* 2012), whilst those in the sub-tropics flower as little as once per year (Luo *et al.* 2017) and desert dwelling *Yucca* species may not flower for several years at a time (Pellmyr 2003). In many tropical fig species, individual trees flower asynchronously throughout the year, resulting in continuous year round flowering at the population level (Bronstein *et al.* 1990; Pereira *et al.* 2007; Peng *et al.* 2010; Chiang *et al.* 2018). The continuous flowering of fig trees is critical to prevent local fig wasp extinction, as constant supply of syconia is required to maintain stable populations (Bronstein *et al.* 1990; Jia *et al.* 2007; Chiang *et al.* 2018). As such, an important question is, how are pollinator populations maintained in OPMs where flowering occurs in discrete episodes and not continuously throughout the year?

Populations of pollinators in OPMs are rarely surveyed (Bronstein *et al.* 1990). The yearly cycle of activity in *Epicephala* moths is especially poorly known. This is probably because of such work requires frequent and long-term observational studies, as well as the inherent difficulties in observing small nocturnal insects. From the few available observations, it would seem that *Epicephala* abundance peaks following periods of host plant fruiting (Kawakita and Kato 2004; Zhang *et al.* 2012; Luo *et al.* 2017). This makes intuitive sense, given that *Epicephala* develop by feeding on growing fruits.

Moths that pollinate plants with discrete and seasonal flowering and fruiting times cannot rely on continuous supply of flowers to maintain their population. As such, it is likely that they may have evolved mechanisms to deal with large gaps in time between fruiting and flowering. Many moths, including at least one species of *Epicephala,* are known to utilise periods of diapause at the egg or pre-pupal stages (Denlinger 1986; Kemp 2001; Sands and New 2008; Luo *et al.* 2017). It seems likely, therefore, that other *Epicephala* moths may utilise some form of diapause during these flowering-fruiting gaps. If diapause does occur in *Epicephala* moths, then we should expect that it should be induced and broken by the same environmental factors that influence flowering. This is because many species of Lepidoptera are known to be phenologically synchronised with their host plants through climatic factors, like temperature (Leland Russell and Louda 2004; van Asch and Visser 2007; Phillimore *et al.* 2012; Fuentealba *et al.* 2017; Posledovich *et al.* 2018). In the *Yucca*-Yucca moth OPM, moths can remain in pre-pupal diapause for up to four years (Pellmyr 2003). An as yet unidentified cue to triggers Yucca moths to emerge at or near the time of flowering. To date there have been no studies of how *Epicephala* moths synchronise their lifecycle with that of their host plant, or the environmental factors influence these interactions.

We set out to determine the annual activity of *Breynia oblongifolia* and it’s *Epicephala* moth pollinators (Finch *et al.* 2018, 2019). *Breynia* is generally regarded to flower and fruit throughout the austral spring, summer and autumn (September to May), meaning that *Epicephala* moths are likely to experience a lack of available flowers during the winter months. However, it is unknown how these *Epicephala* moth populations persist through periods of time in which flowers are absent. We surveyed both host plant and pollinators over multiple years to determine their annual phenology. We hypothesized that because of the obligate dependence of *Epicephala* moths on *Breynia* fruits and flowers, moth abundance would closely track flowering phenology. We also modelled the climatic factors that drive plant and pollinator phenology, predicting that the abundance of adult moths and flowers would be best predicted by the same climatic variables. Then, using our newly gained knowledge of moth and host plant phenology, we looked for evidence of diapause at both the egg and pre-pupal stages. In undertaking this study, we sought to understand how highly specific pollinators track host plant phenology and maintain their populations over time.

## Methods and Materials

### Study species

*Breynia oblongifolia* occurs along the east coast of Australia from southern New South Wales to northern Queensland (Atlas of Living Australia 2018). Mature plants are generally 1-3 m tall and bear unisexual male and female flowers that emerge from the leaf axils (Fig. 1). In male flowers, the stamens are almost entirely enclosed by fused sepals, with only a narrow aperture at the apex allowing access to pollinators. *Breynia* is known to be pollinated by at least two species of *Epicephala* moth, whose larvae develop by consuming around half the seeds in fruits pollinated by female *Epicephala* (Finch *et al.* 2018, 2019). *Epicephala* moths emerge from mature fruits as larvae and pupate on the foliage or leaf litter. However, the annual lifecycle of these highly specific pollinators is largely unknown.

**Figure 1.**
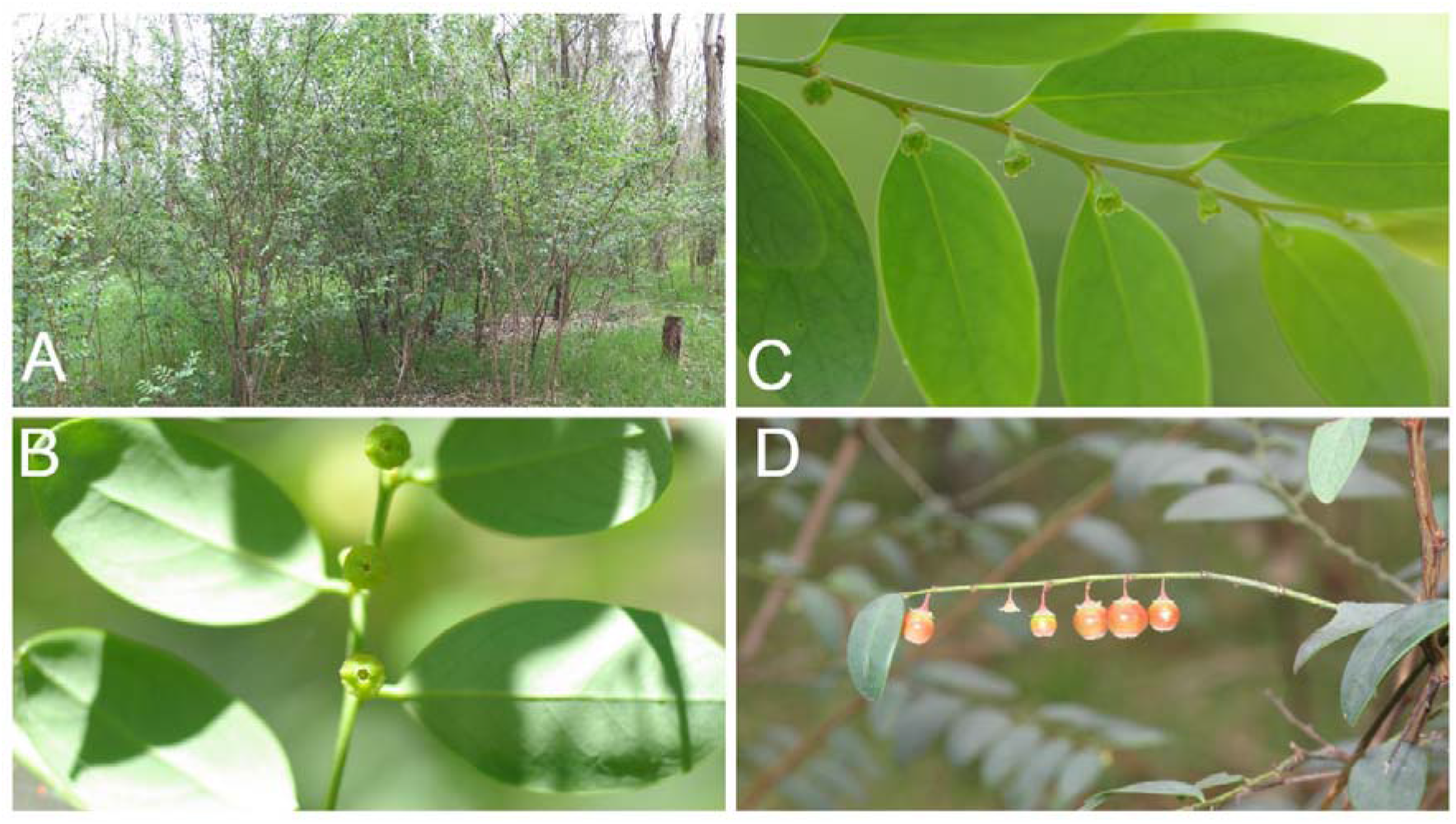
*Breynia oblongifolia*. Mature plants A) male flowers B) female flowers C) female flowers D) mature and developing fruits

### Flowering & fruiting phenology

To determine the occurrence and climatic drivers of fruiting and flowering, we recorded the reproductive phenology of four populations of *Breynia* at two coastal and two inland populations in NSW, Australia. To ensure the independence of climatic effects between sites, we chose four populations that were at least 90 km apart (Table 1). All four populations were situated in *Eucalyptus*-dominated woodlands, where *Breynia* occurs as a component of the understory and had at least 50 flowering plants. Populations at Richmond were surveyed every 2-4 weeks between September 2015 and April 2018. Other populations were surveyed at intervals of approximately 30 days between November 2016 and November 2017. At each site 20 plants were chosen by walking along a pre-existing transect line, either a road or footpath, and choosing the nearest plant >1m in height every 10 m. Four branches, approximately 30cm in length, were selected on each plant and marked for monitoring by repeat survey. On each subsequent visit, the number of male flowers, female flowers, developing fruits (<5mm diameter) and mature fruits (>5mm diameter) were counted on each branch (Fig. 1). In total we conducted 70 surveys of flowering phenology across the four sites; 34 at Richmond and 12 at each of the other three sites. We assessed a total of 100,038 female flowers, 32,473 male flowers, 13,889 developing flowers and 12,098 mature fruits.

**Table 1.**
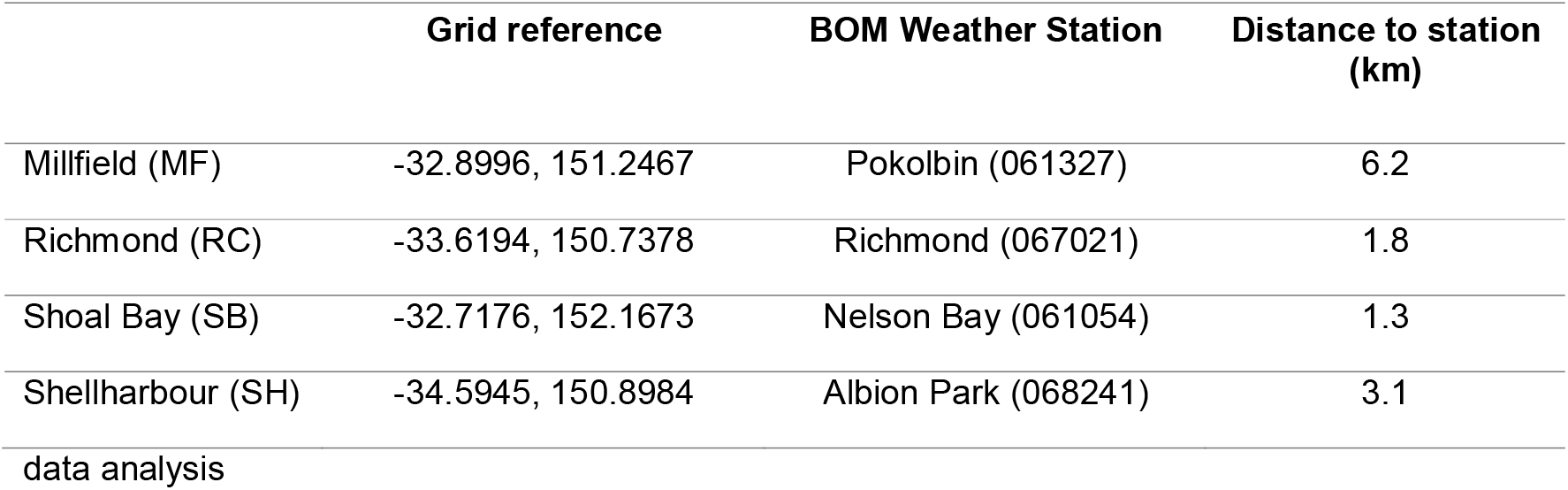
Sampling locations and Australian Bureau of Meteorology (BOM) weather stations used for data analysis

|  | Grid reference | BOM Weather Station | Distance to station (km) |
| --- | --- | --- | --- |
| Millfield (MF) | -32.8996, 151.2467 | Pokolbin (061327) | 6.2 |
| Richmond (RC) | -33.6194, 150.7378 | Richmond (067021) | 1.8 |
| Shoal Bay (SB) | -32.7176, 152.1673 | Nelson Bay (061054) | 1.3 |
| Shellharbour (SH) | -34.5945, 150.8984 | Albion Park (068241) | 3.1 |

All statistical analyses were conducted in R Studio (v1.0.153), using R (V. 1.1.414). Significance was set at α < 0.05. We constructed a series of climatic variables from the climate data to test for associations with flowering and fruiting. We obtained temperature and rainfall data from the Australian Bureau of Meteorology (Bureau of Meteorology, 2018) using the closest available weather station (<7km) at each sampling location (Table 1). Climatic variables included the sum of precipitation in the previous 14, 15-28 and 28-42 days and the sum of all rainfall across all 42 days prior to each phenology observation. In addition, we calculated the average daily minimum and maximum temperatures over the 14 days prior to each phenology observation. Photoperiod is an important climatic factor for the phenology of many Australian plants (Davies 1976; Friedel *et al.* 1993, 1994). As such we chose to include it in our analysis as the duration of daylight hours on the day of observation, to the nearest minute, which we obtained from www.timeanddate.com for Sydney, NSW.

We used negative binomial generalised linear mixed models (GLMM) to determine the degree to which climatic variables influence flowering phenology, whilst controlling for the effects of site and plant. Negative binomial models are useful for modelling count data, such as ours, where a large proportion of “true” zero values results in high variance and over-dispersion (Warton 2005). To determine the most parsimonious combinations of climatic variables on the number of male flowers, female flowers, developing fruits and mature fruits we used the “dredge” function in the MuMIn library to perform model selection. For each reproductive structure (i.e. fruits and flowers), we created a negative binomial model containing all the climatic independent variables. Models were fitted using maximum likelihood and the model with the lowest AICc score for each reproductive structure was then selected for GLMM analysis. We constructed separate negative binomial GLMMs for the number of male, female, developing and mature fruits using the “glmer.nb” function in Lme4 library (v1.1-15) (Bates *et al.* 2015), specifying a random intercept and slope for each site and plant. Where the most parsimonious model identified by dredge included climatic variables that were likely to be highly correlated (i.e. maximum and minimum average daily temperature), we created separate models for each variable and compared them using likelihood ratio tests (Pinheiro & Bates, 2000; Bolker et al., 2009), preferring the model with the lowest AICc score. To check our assumption of over-dispersion we created an alternative GLMM using Poisson regression that does not have an extra parameter for modelling over-dispersion, and compared the two models using a likelihood ratio test. For all reproductive structures (e.g. flowers and fruits), negative binomial regression models performed significantly better than poison regression models (p < 0.0001) supporting our assumption of over-dispersion in the data.

### Pollinator phenology

We conducted surveys of *Epicephala* moths between one and four times per week at the Richmond field site from 13/11/2017 to 01/04/2018. Multiple surveys per week were conducted during periods of high *Epicephala* activity (>4 individuals observed on the first night of observation), otherwise surveys occurred once per week. *Epicephala* moth abundance was observed shortly after sunset when the moths became active. Surveys were conducted by continuously walking along two perpendicular boardwalks to conduct an “X” shaped transect approximately 100 m in total length within the woodland. During surveys we counted the number of *Epicephala* moths observed on plants over a period of 1 hour using a white LED head torch. Although white lights were found to disturb pollination behaviours (Finch *et al.* 2018), using them gave a larger field of view and better chances of detecting *Epicephala* than red lights. No attempt was made to determine which *Epicephala* species were present as this requires destructive abdominal dissections (Finch *et al.* 2018).

We used negative binomial regression to model the abundance of *Epicephala* moth species at the Richmond site over time. To do this, we calculated the mean number of *Epicephala* observed per night for each week by averaging counts of *Epicephala* across all the observations for each week (1-4 observations) from 13/11/2017 to 01/04/2018. For each week, we estimated the sum of male flowers, female flowers, developing fruits and mature fruits at the time of *Epicephala* moth observations using our phenological data. Pearson’s correlation coefficient tests showed that the mean abundance of *Epicephala* moths was significantly correlated with the sum of the number of male flowers, female flowers, developing fruits and mature fruits at the time of each pollinator observation (all |p| < 0.05). As such, we initially modelled *Epicephala* abundance as a function of the sum of male flowers, female flowers, developing and mature fruits using negative binomial regression in MASS package (Venables *et al.* 2002). Although all four phenological variables correlated with *Epicephala* abundance, only the number of mature fruits was found to have a significant relationship (p < 0.05) with the mean number of pollinators observed. The sum of male flowers, female flowers and developing fruits was thereafter excluded from our model.

Because of the significant relationship between *Epicephala* moths and mature fruits (see above), we extended our model of *Epicephala* abundance by adding the variables that drive fruit production, specifically rainfall, temperature and photoperiod. To do this we constructed a range of climatic variables to include in our extended model for each weekly average count of *Epicephala* moths. For rainfall we calculated the total rainfall (mm) in cumulative weekly intervals (0-7, 7-14, 14-21, 21-28 and 0-28 days) prior to each week of observations. For temperature, we calculated the sum of the minimum and maximum daily temperatures, over the same intervals as rainfall, prior to each week of observations. As photoperiod was found to be a significant predictor of the number of female flowers and developing and mature fruits, we also looked for associations between photoperiod and *Epicephala* abundance. Climatic data were again obtained from the Australian Bureau of Meteorology (Bureau of Meteorology, 2018) using the closest available weather station.

To determine the most appropriate timescale for the climatic variables (i.e. 0-7, 7-14 days) we used the “dredge” function in the MuMIn library to perform model selection. This involved creating three separate negative binomial models that contained all the possible variants of rainfall, minimum temperature and maximum temperature. We assumed that the temperature and photoperiod independent variables were likely to be strongly correlated and included photoperiod as a variant of both minimum and maximum temperature in our three models. We used “dredge” again to identify the variable in each model with the lowest AICc run for inclusion in our final model. Because of the large variance in our data and high frequency of zero values, we again opted to use negative binomial regression. To check our assumption of over-dispersion in the data we created an alternative model using the same climatic variables and specifying a Poisson regression model, which does not have an extra parameter for modelling over-dispersion, and compared the fit of the two models using a likelihood ratio test. The log likelihood of the alternative Poisson regression model was significantly lower than the negative binomial regression model (χ^2^ = 392.80, p = 0.007), again supporting our assumption of over-dispersion in the moth abundance data.

### Winter flower surveys

The results of our phenology surveys showed that female flowers were retained over the winter period and began developing to fruits before the appearance of male flowers or moth pollinators (Fig. 4: July-August 2017). As such, we were interested in determining if the female flowers present on *Breynia* over the winter period had previously been pollinated. In the late winter of 2017, we collected ten female flowers from each of 15 *B. oblongifolia* bushes at Richmond. Flowers were selected randomly, from separate branches on each plant. Flowers were taken back to the laboratory and dissected under a Leica EZ4 Stereo Microscope (Leica Microsystems, Wetzlar, Germany) to determine the number of pollen grains per flower and if any insects were present in the flowers. We used a Welch’s t-test to determine if there was a difference in the number of pollen grains in flowers with and without larval feeding damage. We tested for a difference between plants in the number of pollen grains per flower using a one-way ANOVA.

### Pre-pupal diapause

The results of our winter flower surveys suggested that *Epicephala* likely diapause as small eggs or young larvae in female *Breynia* flowers. However, we were also interested in determining if *Epicephala* moths may also diapause between the final larval instar and adult life history stages, as seen in *Yucca* moths. We collected 263 fruits from two successive crops at the Richmond site in the spring of 2018 (November) (n = 115) and the autumn of 2019 (April) (n=148). One to ten fruits were collected haphazardly from ~15 adult *Breynia oblongifolia* plants. Fruits were placed individually in plastic emergence pots (Finch *et al.* 2019). The pots were left outside in a shaded position to mimic natural conditions and checked at one-to-two-week intervals. Insects within the fruits were allowed to emerge naturally. The type of insect, date of larval emergence and date of adult eclosion was recorded for all insects that emerged from the collected fruits. Where no insects emerged within ten weeks of collection, fruits were dissected and checked to see if they had or had not previously contained insects.

## Results

### Flowering & fruiting phenology

Female flowers were generally present throughout the year. Female flowers peaked in abundance during the spring or late summer but declined during winter to low but stable levels (Fig. 2). Interestingly, female flowers were retained on the plants even under periods of drought. For example, in the late winter and early spring of 2017 (July - September) *Breynia* plants at the Richmond site experienced a prolonged spell of unusually dry weather (Fig. S1). During this period, the whole population showed signs of water stress including leaf rolling and leaf abscission. However, many of the plants retained a large proportion of their female flowers, even when they were heavily defoliated. Both the sum of precipitation over the past 42 days (p < 0.0001) and photoperiod (p < 0.0001) were the best predictors of the number of female flowers (Table 3).

**Figure 2.**
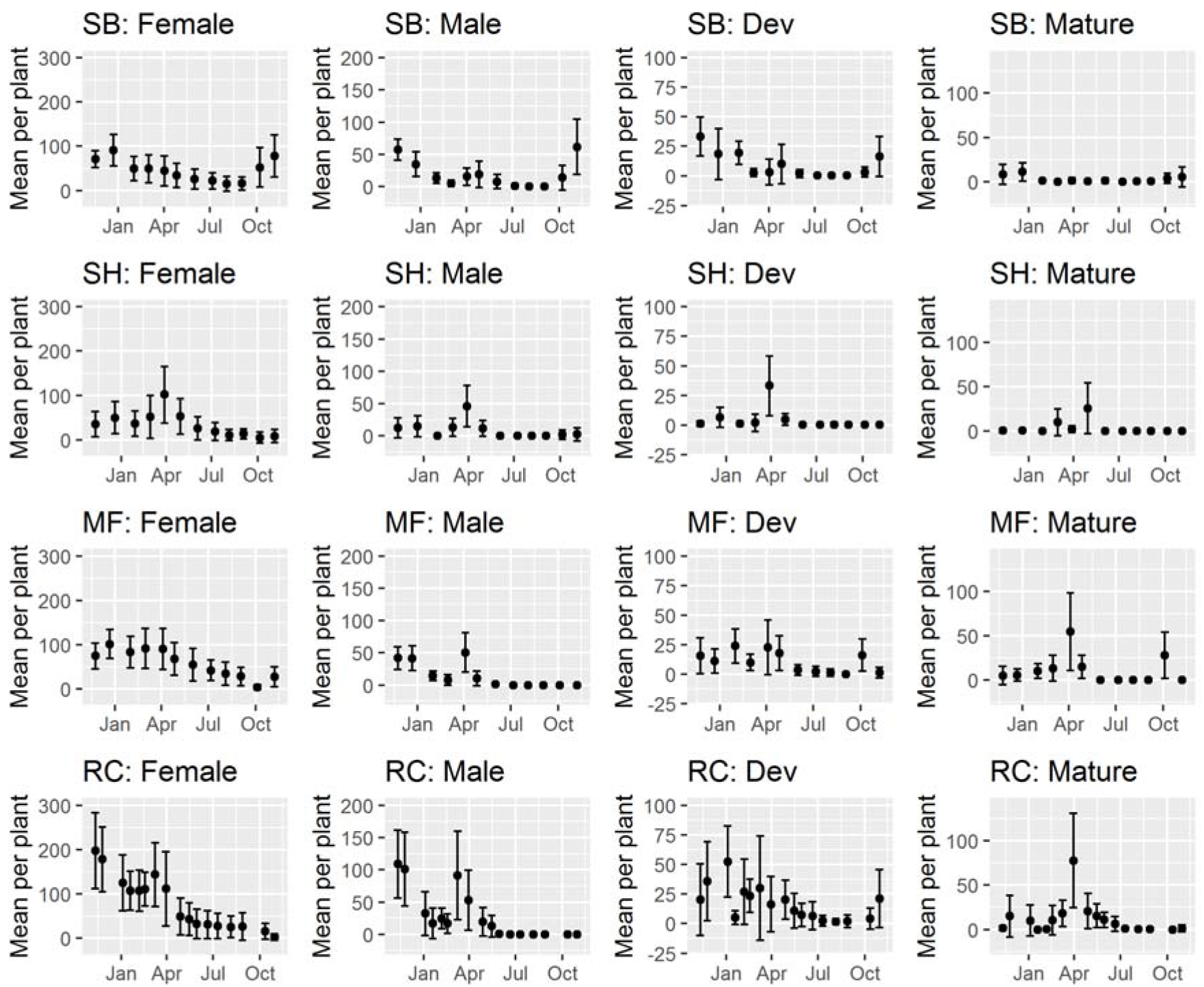
*Breynia oblongifolia* phenology by site. Mean number of male flowers, female flowers, developing fruits (Dev) and mature fruits per plant at Shoal Bay (SB), Shellharbour (SH), Millfield (MF) and Richmond (RC). Error bars show the standard deviation of the mean.

**Figure 3.**
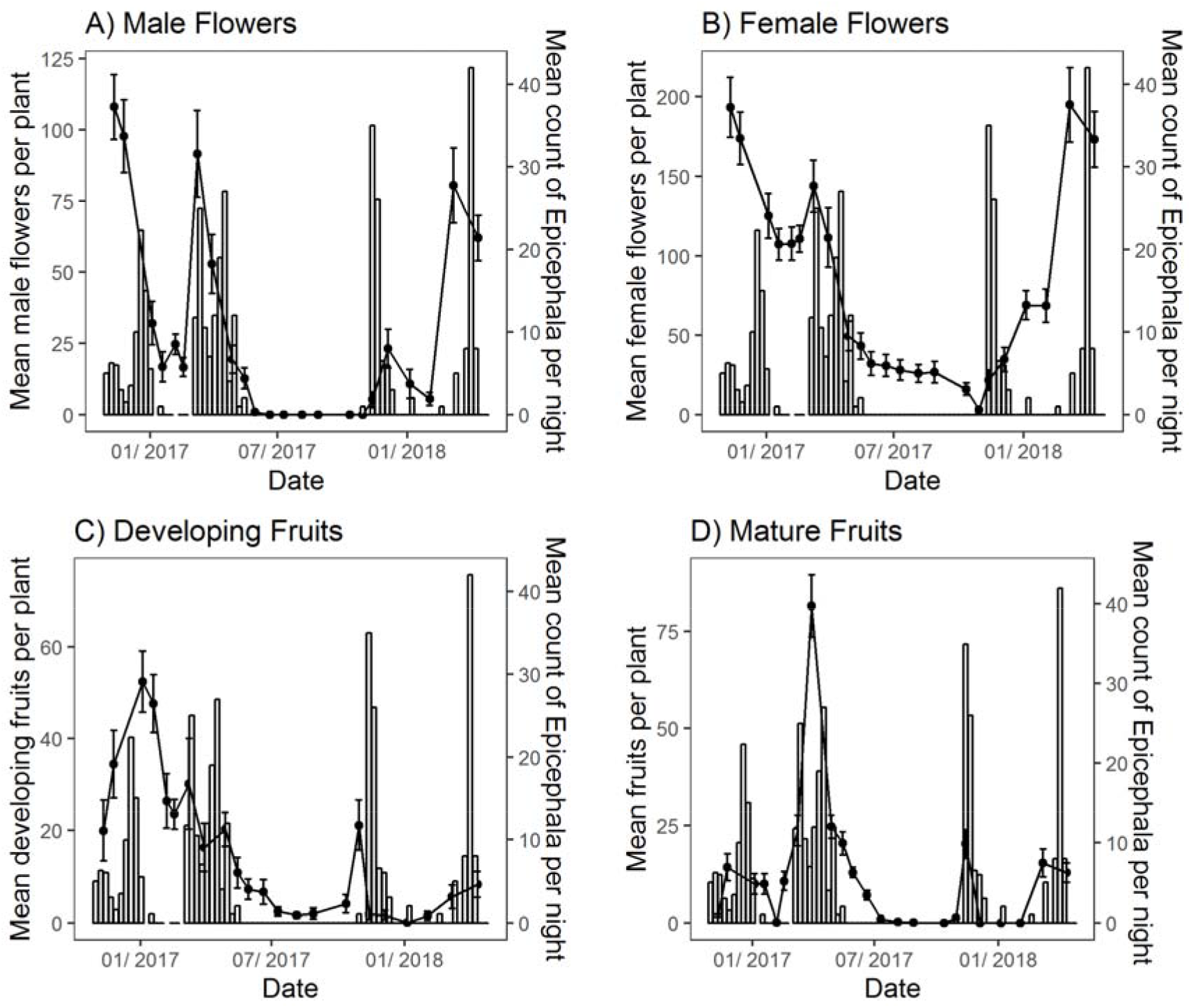
Mean weekly counts of *Epicephala* per night (bars) alongside the mean number of male flowers B) female flowers C) developing fruits D) mature fruits per plant (lines ± SEM) at the Richmond site.

**Table 3.** GLMM coefficients for relationships between climatic variables and flowering and fruiting phenology in our minimal models.

| | $\beta$ | *SE* | *z* | *p* |
| --- | --- | --- | --- | --- |
| **Female Flowers** |  |  |  |  |
| Intercept | 3.7 | 0.182 | 20.47 | <0.0001 |
| Rainfall* | 0.23 | 0.029 | 7.97 | <0.0001 |
| Daylight hours** | 0.58 | 0.027 | 21.22 | <0.0001 |
| **Male Flowers** |  |  |  |  |
| Intercept | 2.06 | 0.242 | 8.52 | <0.0001 |
| Rainfall* | 0.70 | 0.071 | 9.88 | <0.0001 |
| Minimum Temp*** | 1.27 | 0.061 | 20.58 | <0.0001 |
| **Developing Fruits** |  |  |  |  |
| Intercept | 1.65 | 0.33 | 4.88 | <0.0001 |
| Rainfall* | 0.50 | 0.050 | 10.08 | <0.0001 |
| Daylight hours** | 0.60 | 0.045 | 13.41 | <0.0001 |
| **Mature Fruits** |  |  |  |  |
| Intercept | 1.73 | 0.289 | 5.99 | <0.0001 |
| Rainfall* | 0.65 | 0.08 | 7.33 | <0.0001 |
| Daylight hours** | 0.16 | 0.067 | 2.46 | 0.0136 |

\*sum of precipitation over the past 42 days
\*\* photoperiod day on the day of observation
\*\*\*sum of daily minimum temperature over the 14-21 days prior to each phenology observation

Male flowers were most abundant during the spring and summer but declined rapidly in periods of particularly hot and dry weather (Jan-Feb) and were entirely absent during the winter months (May-August) (Fig 2 & 4). As with female flowers, the sum of precipitation over the past 42 days was a strong predictor of the number of male flowers (p < 0.0001). However, in contrast to female flowers, the average minimum temperature (p < 0.0001) was the best predictor of the number of male flowers across the four sites (Table 3).

*Breynia* plants usually produced 1-2 fruit crops per year and each crop lasted for 1-3 months. When combining across all sites, developing and mature fruits were present on plants throughout most of the year (July to May: Fig. 2) but there was considerable variation in fruit production between sites and individual plants, even at the same time of year (Fig. 2). This is most likely because of differences in rainfall that varied between the four sites (Fig. Fruit production did not usually occur during the hottest (Jan-Feb) and coldest parts of year (June and July). However, fruiting was observed to occur in early July at the Richmond site (Fig. 4). Interestingly, at this point in time *Epicephala* moths were not observed to be active (Fig. 4) and male flowers were also absent. Like female flowers, the number of developing fruits was best predicted by the sum of precipitation over the past 42 days (p < 0.0001) and photoperiod (p < 0.0001) (Table 3). Similarly, the abundance of mature fruits was best predicted by the sum of the precipitation over the past 42 days (p < 0.0001) and photoperiod (p = 0.0136) (Table 3).

**Figure 4.**
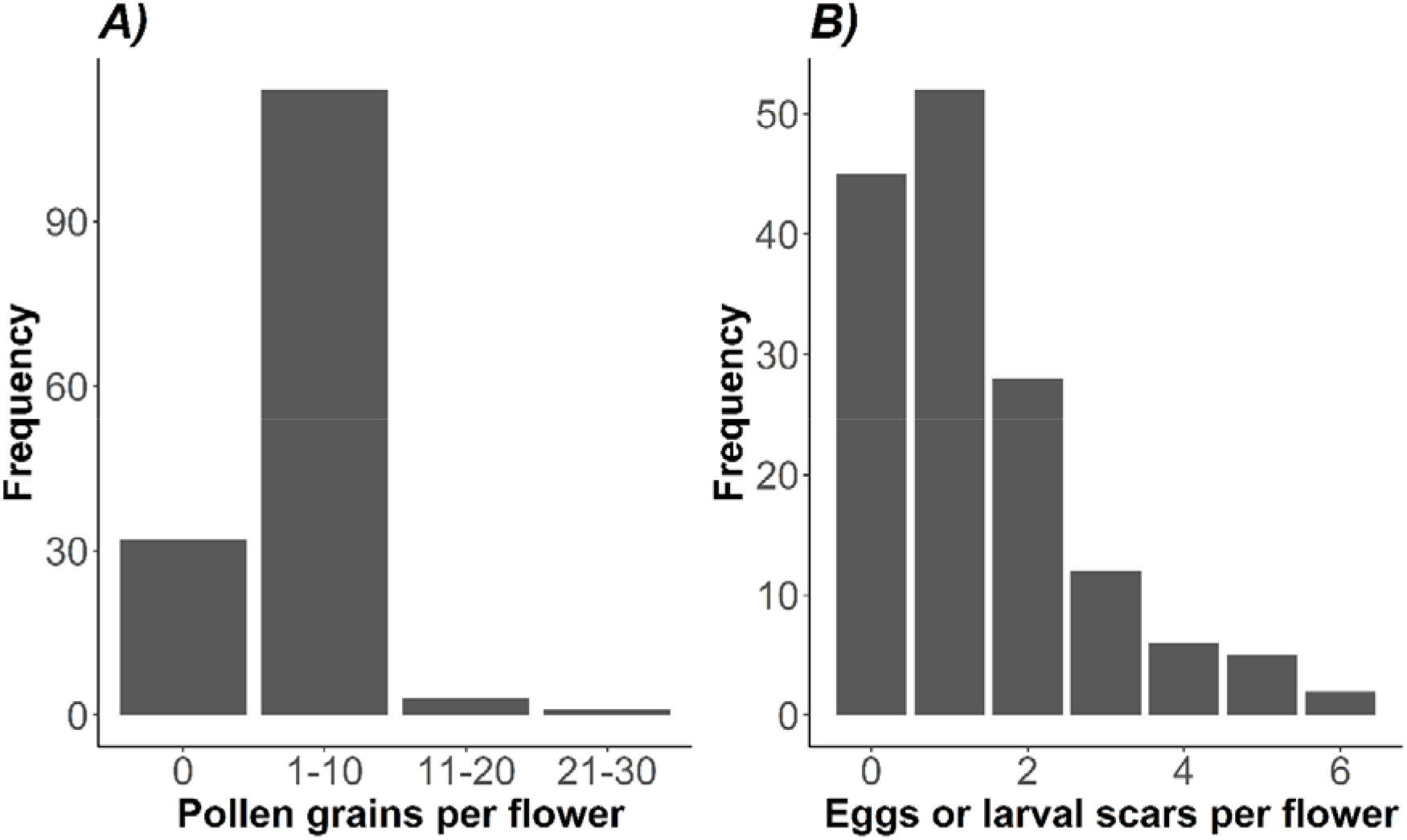
Frequency distributions of A) pollen grains and B) eggs or larval scaring in the overwintering female flowers of *Breynia oblongifolia* collected during in winter (July 2017)(n = 150).

### Pollinator phenology

*Epicephala* moth abundance at the Richmond site varied widely over our 75-week observation period. Adults were seen to be active at the site for periods of 1-2 months in the spring-early summer and then again in the late summer-autumn period but were absent or undetected during the winter months (Fig. 4). In both 2017 and 2018, *Epicephala* were scarcely recorded for up to 3 months during the mid-summer but quickly became very abundant again in the late summer, with up to 42 individuals being recorded in a single hour-long observation period. The mean abundance of *Epicephala* per night was found to correlate significantly with male flowers (0.35, p = 0.0018), female flowers (0.31, p = 0.006), developing fruits (0.25, p = 0.028) and mature fruits (0.53, p < 0.0001). Although fruits and flowers correlated with *Epicephala* abundance, only the number of mature fruits was found to have a significant relationship with the mean number of pollinators observed (Table 2) (Fig. 4). Both the total number of mature fruits (p < 0.01) and the sum of minimum daily temperatures 14-21 days prior to each observation (p < 0.01) were significantly related to the abundance of *Epicephala* adults (Table 2). Rainfall was, however, not related to moth abundance (p > 0.05).

**Table 2.**
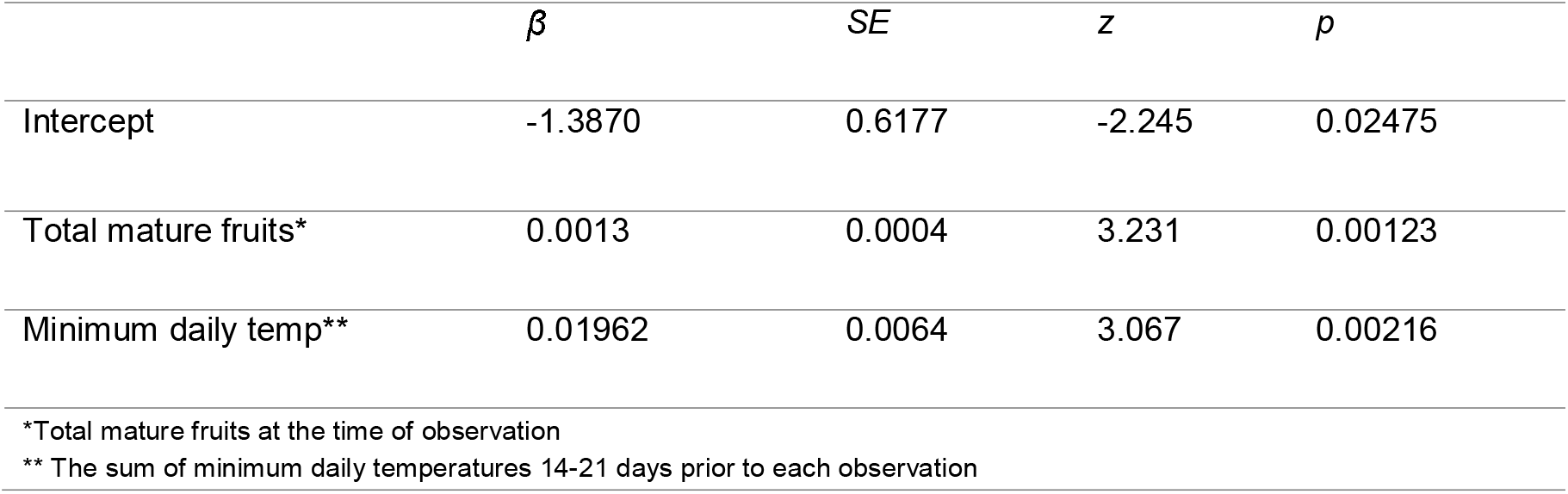
Coefficients for relationships between the number of mature fruits and temperature on the average abundance of adult *Epicephala* per week

| | $\beta$ | $SE$ | $z$ | $p$ |
| --- | --- | --- | --- | --- |
| Intercept | -1.3870 | 0.6177 | -2.245 | 0.02475 |
| Total mature fruits* | 0.0013 | 0.0004 | 3.231 | 0.00123 |
| Minimum daily temp** | 0.01962 | 0.0064 | 3.067 | 0.00216 |

\*Total mature fruits at the time of observation
\*\* The sum of minimum daily temperatures 14-21 days prior to each observation

### Winter flower surveys

Many overwintering female flowers contained pollen and evidence of the presence of overwintering pollinators (Fig. 5). Of 150 inspected flowers, 118 had pollen grains on the surface of the stigma with an average of 10.9 (sd = 5.68) grains per flower. There was no significant difference between plants in the number of pollen grains per flower (*F*_1,148_ = 3.75, *p* > 0.05). Of the 150 flowers collected, 86 showed scars in the tissue of the ovary wall consistent with boring damage by *Epicephala* (Finch *et al.* 2019) and 47 contained at least one *Epicephala* egg or egg case (Fig. 5). These scarred flowers had between 1 and 4 scars (mean = 1.54, sd = 0.78) visible per flower. It was not possible to determine if these marks were attributable to multiple individuals or multiple boring attempts by the same individual. There was no difference in the number of pollen grains present in flowers with or without larval scarring (t = −0.64, df = 130.9, p < 0.520).

**Figure 5.**
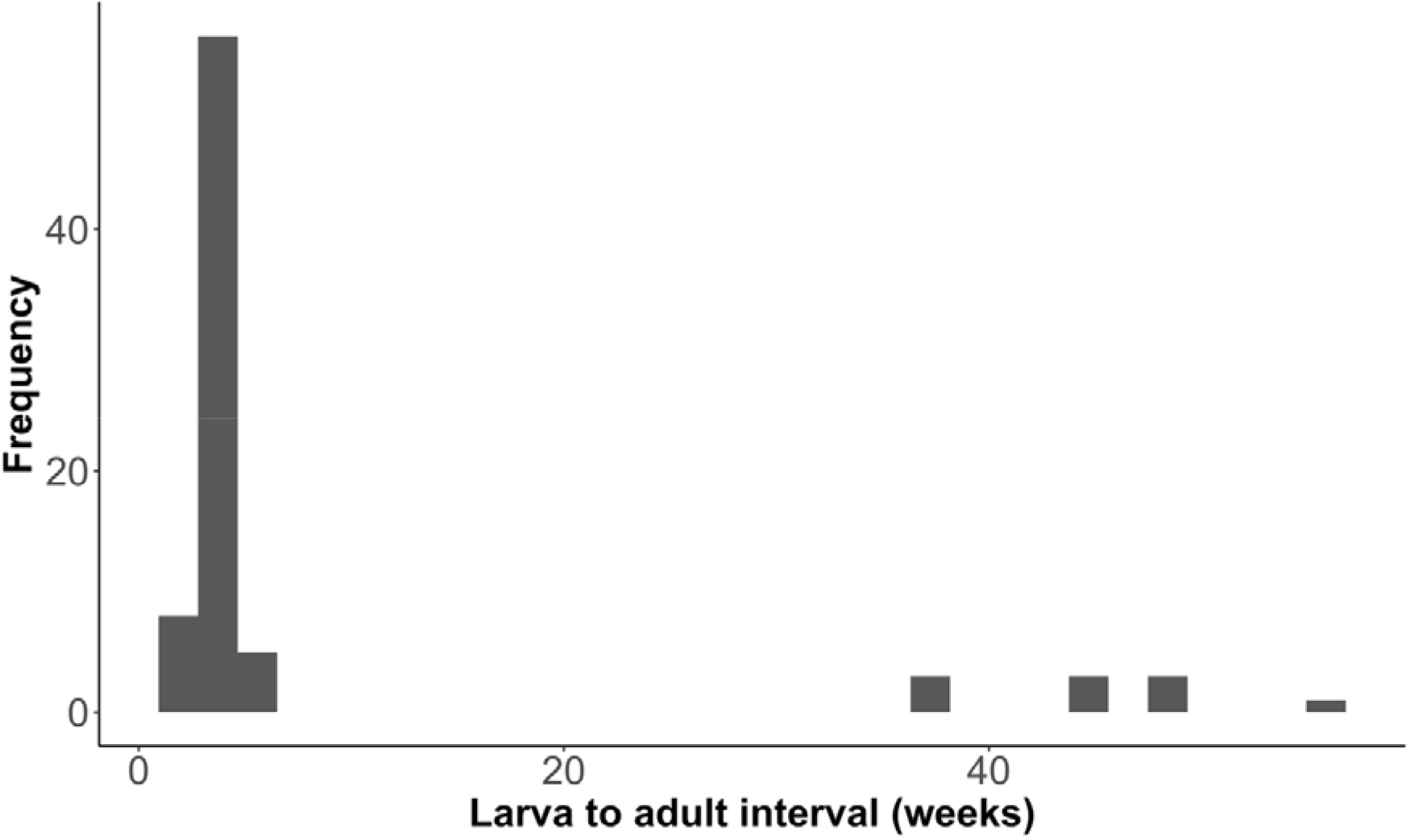
Frequency distribution of the number of weeks between the emergence of *Epicephala* larvae and their eclosion to adults (n = 78).

### Pre-pupal diapause

Of a total of 263 collected fruits, 129 contained no insects, 4 contained non-*Epicephala* insects, 52 contained evidence of feeding damage but no insects (i.e. insects emerged before fruit collection) and 78 contained *Epicephala* larvae. Of the 78 *Epicephala* larvae that emerged from fruits, 68 (87%) pupated and eclosed as adults within 5 weeks with a mean time of emergence to adult eclosion of 3.6 weeks. Ten *Epicephala* (13%) remained as larvae for between 38-56 weeks, successfully pupating and eclosing to adults from nine months to more than one full year after they emerged from fruits (Fig. 5). All Ten moths that remained as larvae for 38 weeks or more were from the autumn crop. Of *Epicephala* collected in the autumn crop, 55 (92%) pupated within 5 weeks, with the remaining 10 (8%) entering an extended pre-pupal diapause. Only 13 *Epicephala* collected from the spring crop, all of which pupated within 5 weeks of emergence.

## Discussion

Flowering and fruiting in *Breynia oblongifolia* were variable between populations and strongly influenced by local rainfall and photoperiod. *Breynia* produced 1-2 fruit crops per year and fruit abundance was a significant predictor of *Epicephala* abundance. The close association between the abundances of fruits and moths suggested that moths may diapause as eggs or young larvae within pollinated flowers, appearing as adults as fruits mature. We confirmed this by showing that many overwintering flowers contained pollen and evidence of diapausing pollinator larvae, henceforth egg diapause. Our observations suggest egg diapause within dormant flowers serves a critical function of physically linking the pollinator lifecycle to host plant flowering phenology, ensuring that pollinator emergence coincides with flowering. Our observations of adult eclosion times also indicate that around 10% of the Autumn generation of *Epicephala* enter a prepupal diapause, eclosing to adulthood after 38-56, equivalent to moth 1-2 generations. However, what factors influence some *Epicephala* to take this alternative strategy is unclear at this stage. In summary, *Epicephala* moths appear to utilise diapause at multiple stages in their lifecycle in order to deal with variable flowering phenology and unpredictable climates (Fig. 6).

**Figure 6.**
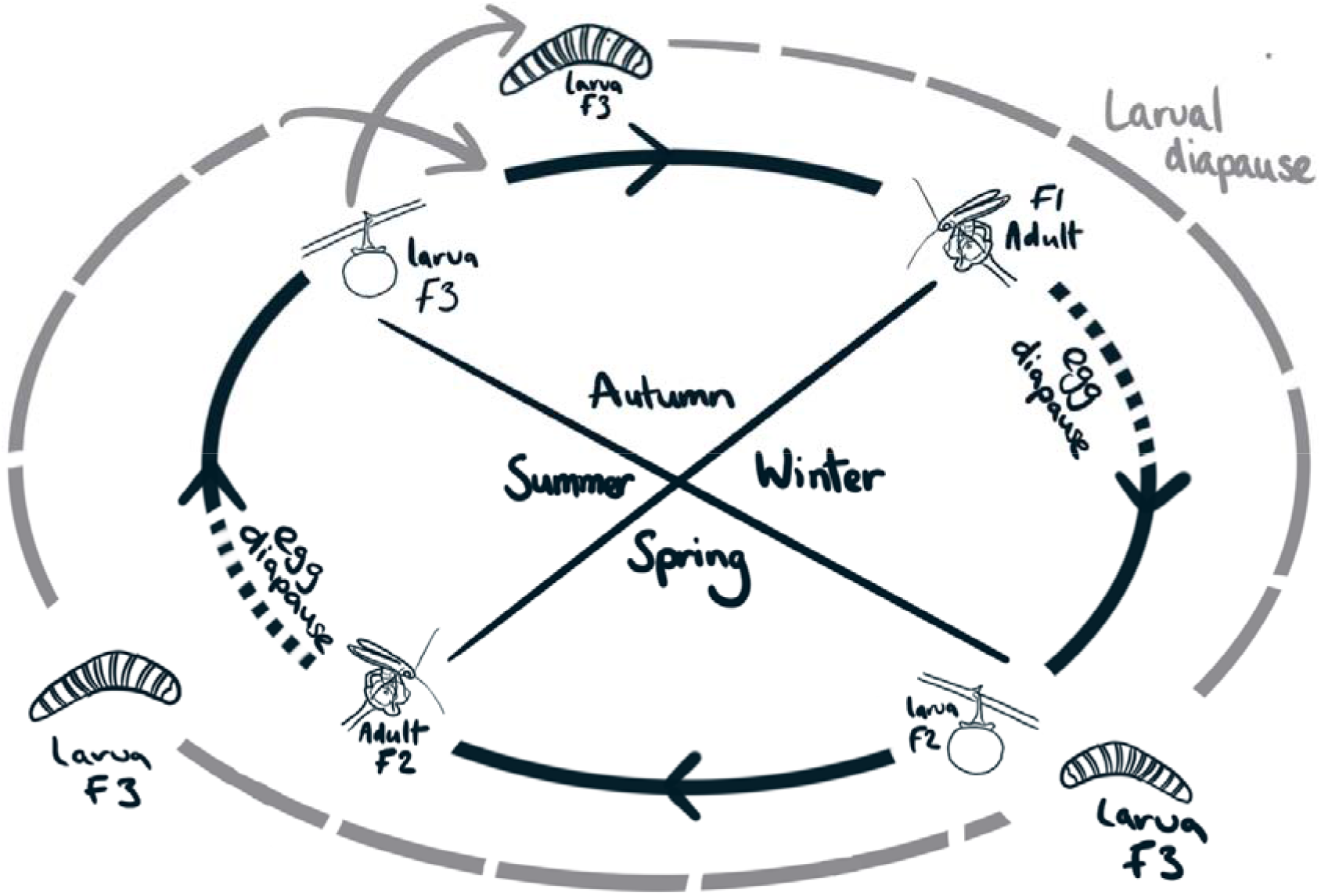
Diagram of the proposed annual lifecycle of *Epicephala* moths on *Breynia oblongifolia* over three successive generations (F1-3). The black cycle denotes the life history for the majority of individuals, in which moths go through two generations per year with the winter months spent as eggs or small larvae in overwintering female flowers. Egg diapause may also occur during the summer months in periods with insufficient rainfall to initiate fruiting. The grey cycle denotes alternative strategy taken by ~10% of individuals in the Autumn crops, in which pre-pupal larvae emerge from fruits and enter diapause for 38-56 weeks, re-joining the general population 1-2 generations later. The number of generations skipped depends on the duration of the diapause. Here, a pre-pupal diapause of 52 weeks is depicted.

### Flowering Phenology

In *Breynia*, Female flowers were present all year round, but their abundance varied widely and was best predicted by rainfall and photoperiod. Inter-annual rainfall can be highly variable within Australia, even when compared to climatically similar areas elsewhere in the world (Nicholls *et al.* 1997; Risbey *et al.* 2009). Much of this variation can be explained by climate-related phenomena such as the El Niño–Southern Oscillation and the Indian Ocean dipole and these events can interact to influence weather patterns across the entire Australian continent (Ashok *et al.* 2003; Wang and Hendon 2007; Risbey *et al.* 2009). As such, we might expect that Australian plants, such as *Breynia*, should favour reproductive strategies that allow them to reproduce at every suitable opportunity (Friedel *et al.* 1994). Indeed, many species of Australian plants are known to fruit and flower in response to both rainfall and photoperiod (Davies 1976; Friedel *et al.* 1994). In this way, *Breynia* seems typical of many Australian plants in that its reproductive phenology is adapted to climatic unpredictability.

The effect of rainfall on a plant’s phenology is likely to be dependent on characteristics of the soil and surrounding landscape, as well as the frequency and intensity of previous rainfall events (Borchert 1994; Jolly and Running 2004; Reynolds *et al.* 2004). In this study, we did not attempt to quantify the effect of soil moisture on phenology because of technical and logistical limitations on collecting such data at multiple field sites over extended time periods. Given that soil moisture more accurately reflects the total water available to each individual plant, we believe that it is likely that soil moisture would perform better than rainfall as a predictor of flowering and fruiting phenology if included in our model.

Interestingly, our winter flower surveys show that *Breynia* frequently retains pollinated flowers over the winter period. Our phenological model found that *Breynia* sets fruit in response to rainfall, which is variable and unpredictable in Australia (Nicholls et al. 1997; Ashok et al. 2003; Dey et al. 2019). Together, these two results suggest that *Breynia* can delay fruit maturation in pollinated flowers over the winter period and potentially also through the spring and summer depending on the prevailing climatic conditions. This likely explains why only female flowers are present during the winter, as well as how some fruits began to develop in the late winter of 2017, when both *Epicephala* moths and male flowers were absent (Fig. 4). This hypothesis would be easily tested with a flower bagging or similar experiment. If confirmed, the observed temporal de-coupling of pollination and fruit development may prove to be an evolved strategy for optimising the timing of fruit production. Retaining previously pollinated flowers that can develop to fruits in response to largely unpredictable rainfall is likely to make *Breynia* less vulnerable to climatic variability, thereby reducing the risk of fruits developing during periods of drought and potential reproductive failure.

The exact mechanism that allows *Breynia* to retain pollinated flowers and then develop them to fruits months later remains unclear. Pollen can remain viable for several hours to several months depending on the species and environmental conditions (Johri and Vasil 1961; Ramsey and Vaughton 1991; Luna et al. 2001; Báez et al. 2002). It may be the that the pollen on the overwintering flowers of *Breynia* have yet to germinate and does so shortly before fruit development. Currently, it is not known how long the pollen of *Breynia* can remain viable under natural environmental conditions. The fruits of several important crop plants are known to arrest their development, with fruit growth initiating again following pollen fertilisation (Wang et al. 2009; Malladi and Klima Johnson 2011). As such, an alternative explanation could be that the retained pollinated flowers are already fertilised but remain in a developmental pause over winter until released by an environmental cue, such as lengthening photoperiod or increased soil moisture following rainfall. Further experimentation is required to answer these questions.

### Pollinator Phenology

*Epicephala* abundance at the Richmond field site in both the spring and summer of 2016-2017 and 2017-2018 occurred in two discrete peaks of high abundance and these peaks coincided with similar peaks in fruit abundance (Fig. 4). Our analysis showed that the number of mature fruits was a significant predictor of *Epicephala* abundance. This is most likely because *Epicephala* emerge from mature fruits as larvae and then pupate to adults (Kato *et al.* 2003; Kawakita and Kato 2004). The appearance of fruits is therefore an important predictor of *Epicephala* moth abundance, as observed in other several other leaf flower species (Kawakita and Kato 2004; Zhang *et al.* 2012; Luo *et al.* 2017). Although we found no statistical evidence for a relationship between rainfall and moth abundance, there is an indirect relationship between the abundance of moths and rainfall via the abundance of fruits that is rainfall dependent. As such, rainfall and photoperiod are critical factors in the phenology of the *Breynia* plant, and by extension its *Epicephala* pollinators.

Consistent with our prediction, both the abundance of male flowers and *Epicephala* moths were best predicted by the mean minimum temperature 14-21 days prior to each plant and pollinator survey. Lepidoptera are often phenologically synchronised with their host plants growth stages through temperature (Leland Russell and Louda 2004; van Asch and Visser 2007; Phillimore *et al.* 2012; Fuentealba *et al.* 2017; Posledovich *et al.* 2018). As male flowers are a source of essential pollen and potentially nectar (Kawakita and Kato 2004; Zhang *et al.* 2012), female *Epicephala* moths are likely to be under selection to synchronize their adult eclosion with the occurrence of male flowers. The fact that both male flowers and moths respond to the mean minimum temperature three weeks prior to each observation probably reflects the developmental time between the onset the environmental stimulus and appearance of mature fruits and moths. Indeed, the average time between the emergence of larvae from fruits and adult eclosion in our eclosion study was approximately three to four weeks. As such, temperature plays a critical role in synchronising moth and pollinator life histories.

The absence of adult moths during the winter months (June-August) indicated that moths diapause during this stage in their lifecycle (Fig. 4). Furthermore, the close association between fruits and the abundance of moths suggested that moths diapause as eggs or young larvae within flowers that then develop to fruit. In line with our expectations, when we examined overwintering flowers, many showed scarring and boring damage consistent with *Epicephala* moths and their larvae (Finch *et al.* 2018, 2019). Periods of egg dormancy are known in *Epicephala* (Luo *et al.* 2017) and many Australian Lepidoptera are also known to exhibit extended periods of egg diapause (Denlinger 1986; Kemp 2001; Sands and New 2008). Our analysis of the phenological data has demonstrated that in *Breynia*, fruiting and flowering are phenologically synchronised with both fruits and new flowers appearing together (Fig. 4). As such, egg diapause likely ensures that overwintering larvae develop and emerge at, or near, the time of flowering. In this way, egg diapause creates a physical link between plant and pollinator lifecycles.

In moths, diapause can occur at both the egg and pre-pupal life history stages. As such, we also checked for diapause at the pre-pupal stage, after *Epicephala* moths had emerged from fruits. Our study of the interval between larval emergence and eclosion found that about 10% of the *Epicephala* larvae that emerged in the autumn entered a pre-pupal diapause for periods of up to 56 weeks (Fig. 5). The majority of moths that emerged in the autumn pupated directly to adulthood, eclosing three to four weeks after leaving the fruits. Spring emerging *Epicephala* were not observed to enter an extended pre-pupal diapause but this may have been because of the relatively low number of *Epicephala* collected (n = 13).

Alternatively, pre-pupal diapause may be exclusive to autumn emerging moths, potentially initiating in response to decreasing temperatures or photoperiod. It would be interesting to see if the proportion of larvae entering pre-pupal diapause remains constant across crops. If the proportion of moths entering into pre-pupal diapause increases in crops maturing later in the season, then pre-pupal diapause may be a facultative response that optimises individual fitness. For example, moths that pupate directly to adults after emerging from late crops may experience shortages of male or female flowers. Conversely, if the proportion of moths using pre-pupal diapause remains constant between crops, then it may function as a bet hedging strategy against future unpredictability in flowering. Bet hedging strategies are common in insects (Hopper 1999) but have not previously been documented in an OPM. Additional collections of *Epicephala* from multiple crops and years will be required to answer this question.

### Implications for the mutualism

Although they are inherently different, it is interesting to compare the phenology of plants involved in the fig, *Yucca* and leaf flower OPMs. Where data is available, it would seem that the flowers of both fig trees and leaf flowers are generally present throughout the year (Bronstein *et al.* 1990; Kawakita and Kato 2004; Pereira *et al.* 2007; Peng *et al.* 2010; Zhang *et al.* 2012; Luo *et al.* 2017). However, as this study and others have demonstrated, flowers may be present but dormant for part of that time. In fig trees, the year round presence of syconia maintains stable fig wasp populations (Bronstein *et al.* 1990). In *Breynia*, and possibly other leaf flower plants, maintaining flowers year-round may also promote stable populations of *Epicephala* moths by providing a refuge for pollinators during periods when plants are not actively flowering. This could also explain why *Breynia* plants at the Richmond site maintained female flowers during a period of obvious drought stress in the unusually dry winter of 2017 (Fig. S1). In the *Yucca*-yucca moth OPM, pollinators diapause as pre-pupal larvae in the soil around their host plants and may wait several years between the appearance of flowers (Pellmyr 2003). The ability of yucca moths to diapause for multiple years between flowering events probably also promotes the stability of pollinator populations. Our study has found that both year-round flower provision and pre-pupal diapause occur in at least some leaf flower plants, showing similarities with both the *Yucca* and fig OPMs. It remains to be seen if these factors promote the stability of pollinator populations in leaf flower plants.

Egg diapause in *Epicephala* may have other important implications for the mutualism. Overwintering larvae or eggs are likely to suffer moderate levels of mortality during diapause (Régnière and Duval 1998; Elverum *et al.* 2003). This increased mortality may benefit the plant by reducing the number of seeds that are consumed by pollinators. Overwintering mortality could help to explain the large number of *Breynia* fruits (10-30%) that do not contain *Epicephala* larvae (Finch *et al.* 2019). Fruits that do not contain pollinators are generally more frequent in crops collected in the spring and contribute a high proportion of the intact seeds produced across *Breynia* populations (Finch *et al.* 2019). As such, egg diapause mortality may be an important factor in reducing seed destruction by pollinating seed herbivores, thereby helping to maintain the mutualism in the face of competing interests between mutualists.

At least one species of *Glochidion* (Phyllanthaceae) is known to delay fruiting for up 11 months after pollination and ovipositon by *Epicephala* moths (Luo *et al.* 2017). As such, it remains to be seen how widespread delayed fruit development, egg diapause and pre-pupal diapause may be within the leaf flowers and leaf flower moths and, furthermore, to what degree it may have facilitated the evolution of these highly diverse mutualisms.

## Supplementary figures

**Figure S1.**
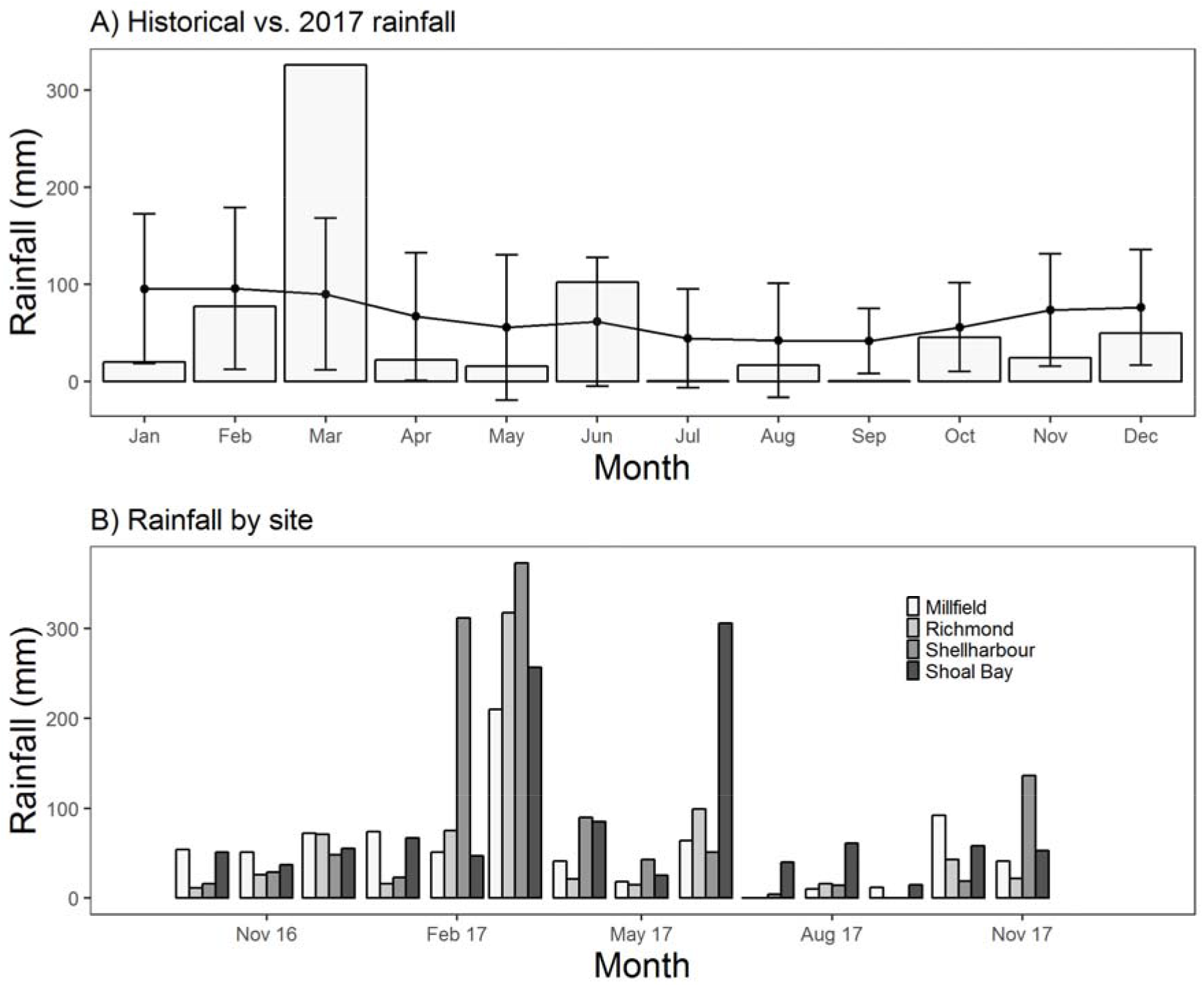
Historical and contemporary rainfall trends A) Total rainfall per month at Richmond during 2017 (Bars) against the average total monthly rainfall at the site since 01/01/1881 (Lines with error bars ± SD) and B) Total monthly rainfall at the four sampling locations from October 2016 to November 2017.

